# HIV-1 drug resistance profiling using amino acid sequence space cartography

**DOI:** 10.1101/2021.07.31.454569

**Authors:** Karina Pikalyova, Alexey Orlov, Arkadii Lin, Olga Tarasova, Gilles Marcou, Dragos Horvath, Vladimir Poroikov, Alexandre Varnek

## Abstract

Human immunodeficiency virus (HIV) drug resistance is a global healthcare issue. The emergence of drug resistance demands treatment adaptation. Computational methods predicting the drug resistance profile from genomic data of HIV isolates are advantageous for monitoring drug resistance in patients. Yet, the currently existing computational methods for drug resistance prediction are either not suitable for complex mutational patterns in emerging HIV strains or lack interpretability of prediction results which is of paramount importance in clinical practice. Hence, to overcome these limitations, new approaches for the HIV drug resistance prediction combining high accuracy and interpretability are required. In this work, a new methodology for the analysis of protein sequence data based on the application of generative topographic mapping was developed and applied for HIV drug resistance profiling. It allowed achieving high accuracy of resistance predictions and intuitive interpretation of prediction results. The developed approach was successfully applied for the prediction of HIV resistance towards protease, reverse-transcriptase and integrase inhibitors and in-depth analysis of HIV resistance-inducing mutation patterns. Hence, it can serve as an efficient and interpretable tool to suggest optimal treatment regimens.

## Introduction

The global epidemic of human immunodeficiency virus (HIV) infection and acquired immunodeficiency syndrome (AIDS) is one of the major public health issues, affecting more than 38 million people worldwide (UNAIDS, 2020). The primary causative agent of HIV infection is the *Human immunodeficiency virus 1* belonging to the *Lentivirus* genus, *Retrovidae* family (Knipe & Howley, 2013). The virus induces a progressive weakening of the immune system and if untreated, leads to a rise of opportunistic infections and, subsequently, death. The development of a preventive vaccine remains a challenge, and, for now, the treatment of the HIV/AIDS is based on the usage of antiretroviral therapy (ART) (Ananworanich, 2015; Pavlakis & Felber, 2018). Since its introduction, ART has a significantly extended life expectancy and improved the quality of life of people diagnosed with HIV infection (Knipe & Howley, 2013). Currently recommended ART consists in the treatment by a combination of inhibitors of HIV enzymes including reverse transcriptase (RT), integrase (IN), and protease (PR) (WHO, 2018). Despite being highly effective in many cases, ART failure can occur for some individuals due to the emergence of resistance against one or more inhibitors (Iyidogan & Anderson, 2014). Drug resistance testing is therefore essential for the selection of an optimal ART regimen (Günthard et al., 2018). The drug resistance can be assessed by either phenotypic or genotypic assays. Phenotypic assays provide information about the inhibitory activity of drugs against a particular virus variant in *in vitro* cell-based experiments. Although these assays provide information essential for establishing genotype-phenotype correlations, they are labor- and time-consuming, which restricts their use in routine clinical diagnostics. The genotyping assays, on the other hand, are based on genome sequencing, which is accessible, fast, and inexpensive, but requires an accurate interpretation system allowing one to predict the resistance profile for the particular virus variant. Due to the rapidity and availability of sequencing, a large amount of sequence data was accumulated in public repositories, and interpretation systems based on various computational methods for predicting drug resistance were developed to assist in clinical diagnostics.

One of the largest and commonly used public HIV drug-resistance databases is the Stanford HIV drug resistance database (HIVDB) (Rhee et al., 2003). It comprises more than 400 thousand sequences of HIV-1 proteins and over 60 thousand drug resistance profiles for RT, PR, and IN proteins. Numerous algorithms were suggested for the interpretation of mutational patterns present in these data and the prediction of the mutations’ influence on drug resistance (Vercauteren & Vandamme, 2006). These algorithms can be broadly classified into rule-based and machine learning (ML)-based. The rule-based approaches comprise a set of pre-defined rules according to which the variant is defined as resistant or susceptible. They are typically derived from published data on mutations associated with drug resistance and correlations between treatment regimen and virological response data (Vercauteren & Vandamme, 2006). Although, rule-based methods are still widely used due to their interpretability, they require a continuous update of the existing rules and are not very accurate when complex mutational patterns are present in sequences (Singh, 2017). Alternatively, ML algorithms can be used to overcome the aforementioned problems of rule-based approaches. Various ML algorithms were applied for HIV drug resistance profiling, including linear regression (Rhee et al., 2006; Yu et al., 2014), support vector machines (Khalid & Sezerman, 2018; Masso & Vaisman, 2013; Rhee et al., 2006), decision trees (Beerenwinkel et al., 2002; Rhee et al., 2006), random forest (Shen et al., 2016; Tarasova et al., 2018), artificial neural networks (Pasomsub et al., 2010; Rhee et al., 2006; Sheik Amamuddy et al., 2017; Steiner et al., 2020; Wang & Larder, 2003), and Bayesian approaches (Tarasova et al., 2017). They allowed achieving better accuracy of drug resistance predictions as compared to rule-based methods.

Despite the high accuracy of resistance predictions, most ML algorithms lack intuitive interpretability inherent to rule-based methods. The ideal algorithm would need to combine high predictive accuracy and intuitive interpretation of the results of the predictions. One way to get an interpretable machine learning system is to combine the visualization of data with predictions that would be consistent with the results of the visualization. To represent the sequence data, dimensionality reduction methods can be used. In such approaches, biological sequences are encoded as objects in a multidimensional space and then projected to 2D or 3D spaces. In recent years, several algorithms dimensionality reduction such as: principal components analysis (PCA) (Hotelling, 1933), self-organizing maps (SOMs) (Kohonen, 1982), generative topographic mapping (GTM) (Bishop et al., 1998), etc. were applied for sequence space analysis. The resulting maps obtained after applying aforementioned dimensionality reduction methods can be “colored” by any property of the items it hosts (e.g. the map hosting PR sequences can be coloured by associated drug resistance profiles). Hence, the “property landscapes” are produced – the local “color” representing the mean of experimental properties of the items residing in the given point or zone of a map. Depending on the specific hypotheses and the pertinence/information-richness of the “training set” of items used to create such landscapes, these can be used as predictive tools – by assuming that the unknown property of a new item can be read from the “color” of the landscape zone in which it resides. Regarding HIV, SOM and kernel PCA were applied for visual analysis of the HIV mutant sequences’ space (Drǎghici & Potter, 2003; Ramon et al., 2019).

Among other dimensionality reduction techniques, GTM seems to be particularly useful, as it has a rather unique ability to be both a visualization tool and a multi-task quantitative predictive model. GTM was extensively studied for chemical space analysis and showed superior visualization ability and prediction accuracy in tasks related to quantitative structure-property relationships modelling as compared to other dimensionality reduction algorithms (Gaspar et al., 2016).

Herein, the GTM method was used for building interpretable maps of HIV sequence space and predicting HIV drug resistance profiles. The accuracy of GTM-based resistance predictions was comparable with the ones of other state-of-the-art ML algorithms: random forest (RF), support vector machines (SVM), and gradient boosting (GB). In addition, we combined the analysis of sequence space maps with algorithms allowing to identify characteristic mutations of sequences clustering in the same parts of the map in order to visualize mutation patterns leading to the resistance. Thus, in this work, an interactive tool for sequence space exploration was developed and applied to the HIV drug resistance analysis. The developed approach enables one to work with high-dimensional genomic and proteomic data, all along by ensuring the rapid building of interpretable models. It does not only overcome the interpretability problem inherent to ML methods but also provides a general methodology applicable for various tasks related to phenotypes prediction. Hence, we believe that this approach will be useful in numerous tasks related to the analysis and exploration of sequence space and modelling of quantitative genotype-phenotype relationships (QGPR).

## Methods

### Data acquisition and pre-processing

The HIV protein sequence data with associated resistance profiles accumulated in large databases provide the basis for computational drug resistance predictions. The resistance is expressed by fold ratio (FR) value:

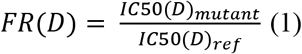

*IC*50(*D*)_*mutant*_ − drug concentration at which the replication of the mutant virus strain is inhibited by 50%, *IC*50(*D*)_*ref*_ − drug concentration at which the replication of highly susceptible reference virus strain is inhibited by 50%.

The above ratio is specific for each anti-HIV drug D, and these drugs were each targeted at a specific HIV protein. The challenge of this work is to predict *FR*(*D*) as a function of the mutations affecting the viral protein against which D was designed for (its primary target). In principle, mutations affecting other viral proteins might potentially indirectly impact on the effectiveness of the drug D, but the analysis of such effects, if ever observed, is beyond the purpose of the current paper. Specific models will thus be built for various drugs binding HIV reverse transcriptase, integrase, and protease sequences. Associated fold ratio values from high quality filtered HIVDB genotype-phenotype dataset were used for modelling quantitative genotype-resistance relationships, where “genotype” in this context designs the (mutant) protein sequences, the various “phenotypes” being *FR*(*D*) for each drug binding to the target. Overall, 1707, 659, and 1958 mutant HIV RT, IN, and PR sequences respectively with corresponding FR values measured for six RT, two IN, or eight PR inhibitors were extracted from the HIVDB genotype-phenotype dataset. Each mutant protein sequence in the database was represented as a set of amino acid substitutions in certain positions in regard to the reference sequences. These mutant protein sequences were then completed by original non-mutated amino acid residues from the consensus sequence to form viral protein chains. The number of amino acid residues was 99 per HIV PR monomer, 288 for the HIV IN, and 240 for the HIV RT. Although the p66 subunit of HIV RT comprises 560 amino acid residues, the filtered data on mutations and associated resistance profiles was available only for the first 240 amino acid residues of the RT p66 domain and, hence, only this data was used for modelling. Since the first three N-terminal amino acids were not systematically reported for all protease sequences, they were ignored altogether. Some of the sequences contained non-standard symbols, such as the “.” symbol denoting unknown amino acid, “X” symbol denoting a mixture of four and more amino acids, symbols denoting the presence of two or more amino acids at the same position (“mixtures of amino acids”, e.g., “A/S” for alanine-serine mixture), etc. The amino acid mixtures arise due to the noise in the experimental data or the coexistence of several HIV variants in a single patient’s virus sample. The presence of these amino acid mixture symbols indicates ambiguities in sequence data and can confound QGPR modelling (Ramon et al., 2019). Several approaches were suggested for the processing of such sequences. In a majority of approaches the sequences with symbols denoting insertions and deletions, unknown amino acids, and mixtures of four and more amino acids (“X” symbol) were deleted (Masso & Vaisman, 2013; Ramon et al., 2019; Yu et al., 2014). By contrast, the strategies for processing the sequences with amino acid mixtures were diverse. Tarasova et al. (Tarasova et al., 2018) addressed this problem by verifying the frequency of occurrence of amino acids from mixtures in the set of corresponding positions in the examined enzymes of other drug-resistant HIV-1 variants and keeping the most frequent one. Masso et al. (Masso & Vaisman, 2013) deleted all the sequences with the mixtures of amino acids. In contrast to the latter approach, the technique leading to the expansion of the dataset via combinatorial enumeration of all possible variants of the sequences containing mixtures was also used (Sheik Amamuddy et al., 2017; Shen et al., 2016; Yu et al., 2014). Ramon et al. (Ramon et al., 2019) suggested applying various kernels for handling amino acid mixtures, hence maintaining the authenticity of the data and reducing the chance of incorrect mutation patterns being introduced.

Taking into account the aforementioned information, all sequences containing symbols, which did not denote 20 standard amino acids were removed from the dataset. HIV IN sequences containing amino acid mixtures located in major drug resistance mutation positions − positions that, according to current knowledge (Rhee et al., 2003), are associated with significant changes in drug resistance (DRM positions), were removed. HIV PR and RT sequences of high quality filtered HIVDB genotype-phenotype dataset did not contain any mixtures in DRM positions. Amino acid mixture symbols not located in major drug resistance mutation positions were replaced by the symbol of the first amino acid from the mixture symbol (i.e., a mixture such as “A/S” was replaced by the “A” symbol). Moreover, only sequences corresponding to the most studied HIV subtype B were kept, to increase the prediction performance for this subtype. Finally, 1581 sequences with known FR for PR inhibitors, 510 sequences with available FR for IN inhibitors, and 1389 sequences with known FR for RT inhibitors were prepared. Classes “resistant” and “susceptible” were assigned to each of these sequences with respect to every drug based on the corresponding FR values and clinical thresholds extracted from HIVDB (Table S1, Supplementary Data). The GTM-based cartography includes as a first step the unsupervised construction of a map spanning the relevant space of possible protein sequences as defined by a pool of representative items (the “frame set”) (Lin et al., 2020). The latter do not need to be annotated by FR – therefore, the frame set was expanded to include also observed mutant sequences for which FR values were not yet reported for the studied drugs. Separate frame sets for RT, IN, and PR sequences were composed from the corresponding labelled sequences and unlabelled sequences from Genotype-Treatment dataset from HIVDB. In such a way, the individual frame sets consisted of 6591 sequences for IN, and 5000 sequences for PR and RT. The enrichment of the dataset with the randomly selected unlabelled sequences was performed to ensure more uniform coverage of the mutant HIV proteins sequence space. Since additional unlabelled sequences are genetically different from the labelled ones, the enriched dataset better delineates the sequence space of HIV mutants.The workflow used for the data pre-processing is shown in Figure S2, Supplementary Data.

### Descriptors

A prerequisite for applying most machine learning algorithms is encoding amino acid sequences into numeric vectors (Zamani & Kremer, 2011). Herein, various types of encoding schemes were used: k-mers (subsequences of amino acids of length k), one-hot encoded vectors derived from BLOSUM, and HIVb amino acid substitution matrices. Reduced amino acid alphabets, combining amino acids by their physico-chemical properties, and allowing to simplify protein complexity (Zheng et al., 2019) were also applied. In addition to that, descriptors transformed with Principal Component Analysis (PCA) and kernel PCA were generated. The scheme of the process used for sequence descriptors generation is shown in Figure S3. Presentation of the descriptors used in this work is given in Text S3, Supplementary Data.

### Generative Topographic Mapping

Generative Topographic Mapping (GTM) is a non-linear dimensionality reduction method that was introduced by C. Bishop et al. (Bishop et al., 1998). The GTM method is based on the injection of a flexible hypersurface (the “manifold”) into the high-dimensional data space formed by the given data points. In the current work, each data point corresponds to an HIV protein sequence encoded by a numerical vector of a fixed length (Figure S3). The manifold is the center of a normal distribution modeling the data distribution; the algorithm optimizes the shape of the manifold and the width of the normal distribution to maximize the likelihood of the dataset. Once the manifold is fitted, the data points are projected onto it. Finally, a 2D map represents the unbent manifold with sequences projected onto it. Note, that GTM considers each sequence not only as a single point but also as a probability distribution in its latent space (i.e., a probability to find the given sequence residing in the given node of the map). This allows one to create a cumulative landscape obtained by overlapping the probability distributions of all the training items (Kireeva et al., 2012). The latter can then be “colored” by associating to each node of the map the mean property/activity class of sequences residing in it, hereby generating fuzzy classification or continuous property landscapes. These landscapes can be used as predictive models at the same time providing data visualization. The GTM algorithm applied to amino acid sequences space is shown in Figure 1.

**Figure 1.**
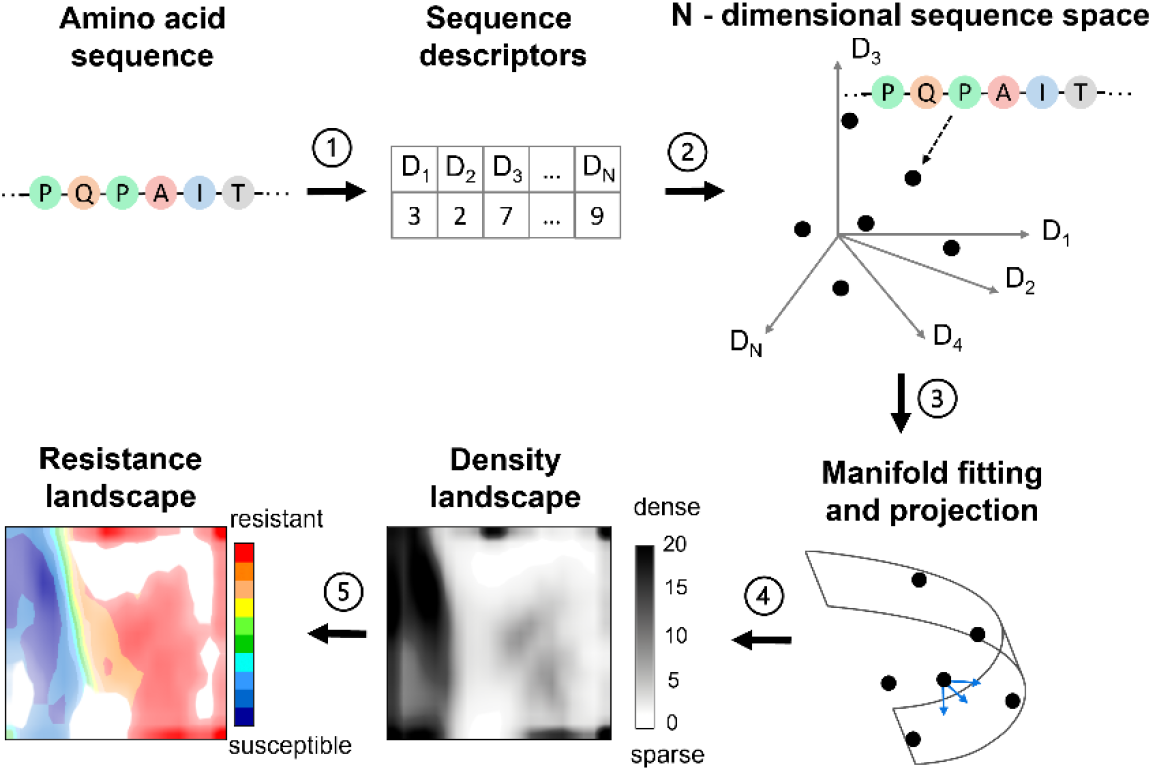
Key steps of the GTM algorithm applied to amino acid sequence space. Each amino acid sequence is encoded by a numerical vector (1) defining its position in the N-dimensional descriptor space (2). The flexible manifold is fitted in a way to approach the data points followed by projection of the data points onto the manifold (3). A 2D map results from the unbending of the manifold. Each projected datapoint is characterized by a probability to be located in the nodes of a rectangular grid superposed with the manifold. (4) Each node is then associated (“coloured”) with a weighted average of resistance values of residing sequences. Ensemble of the coloured nodes forms the resistance landscape (5).

GTM has four hyperparameters: map resolution, number of hidden Radial Basis Functions (RBF), regularization coefficient, and width of an RBF. Along with descriptors type, the best GTM hyperparameters values for the given modelling task must be found. For this purpose, Genetic Algorithm (GA) is applied.

### Genetic algorithm

The GA is a stochastic evolutionary optimization algorithm adapted for parameter optimization problems. In our study, the GA was used as a method for selecting the optimal descriptors and hyperparameters of the GTM (Horvath et al., 2014). To build the GTM manifold, a set of 5000 HIV protein sequences (frame set; see the description above) is used. The selection of appropriate descriptors and the GTM hyperparameters was carried out according to the evaluation of the model’s predictive performance. The latter was estimated by an averaged balanced accuracy (average proportion of sequences predicted correctly for each class: susceptible and resistant) for each drug in a 5-folds cross-validation procedure:

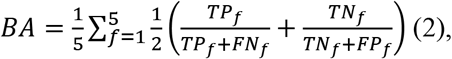

**TP_f_** is the number of truly resistant sequences predicted as resistant in the fold f,

**TN_f_** is the number of truly susceptible sequences predicted as susceptible in the fold f,

**FN_f_** is the number of truly resistant sequences predicted as susceptible in the fold f,

**FP_f_** is the number of truly susceptible sequences predicted as resistant in the fold f.

The best maps are thus the ones which are able to host, on their one manifold, a maximum of highly predictive resistant/susceptible binary classification landscapes (corresponding to the different drugs associated to that protein). Using a common manifold to model drug resistance of various drugs based on a same manifold not only enhances the interpretation (by providing a common reference system to navigate the sequence space) but also presents the inherent benefits of multi-task learning (Lin et al., 2019). However, different protein targets require distinct dedicated maps, each covering a given sequence space (of IN, PR and RT, respectively).

### Comparison to other machine learning methods

The GTM’s drug resistance predictive ability was compared with other state-of-the-art machine-learning algorithms implemented in the sci-kit learn library (v. 0.23.1) (Pedregosa et al., 2011). The following hyperparameters were tuned during optimization (grid search):

- RF (Breiman, 2001): number of trees (100, 300, 500, 1000), number of features (all features, squared root of the number of features, log2 of number of features), out-of-bag sampling (with and without), max depth of trees (full tree, 5, 10, 30), class weight (none, balanced, balanced_subsample);
- SVM (Boser et al., 1992): regularization coefficient (0.1, 1, 10, 100, 1000), kernel coefficient (1, 0.1, 0.01, 0.001, 0.0001), kernel (‘rbf’, ‘linear’, ‘poly’, ‘sigmoid’);
- GB (Friedman, 2001): number of trees (100, 300, 500, 1000), number of features (all features, squared root of the number of features, log2 of number of features), learning rate (0.0001, 0.001, 0.01, 0.1, 1.0), subsampling (0.5, 0.7, 1.0) max depth of trees (full tree, 5, 10, 30).

Evaluation of the model performance was made using 5-fold cross-validation (the same as in the case of GTM).

### Analysis of resistance determining mutation patterns

The combination of mutations in certain amino acid positions (mutation pattern) defines the position of the sequence on the map. Hence, one can extract sequences residing in the particular node or a group of nodes (a zone) and determine which particular mutation pattern is prevalent in these sequences. This analysis can be performed using the algorithms for the identification of specificity determining positions. In this work, the SDPred algorithm v.2 (Kalinina et al., 2009) was used. This algorithm is based on the calculation of the position-wise mutual information in two or more groups of aligned sequences. It allows identifying positions in which the amino acid distribution is significantly different between groups of sequences. The algorithm was applied in combination with the GTM: sequences residing in the specific node or zone of the GTM were considered as the target group and all other sequences projected onto the map in other nodes were considered as another group.

The graphical representation of aligned amino acid sequences (i.e. sequence logo) was performed with Logomaker python library (Tareen & Kinney, 2019). The sequence logo representation is composed from stacks of symbols that reflect the presence of particular amino acids in a specified position in a group of aligned sequences. The height of the amino acid symbols within the specified stack (position in a sequence) represents the frequency of occurrence of a certain amino acid in a group of aligned sequences.

## Results and Discussion

### Cartography of HIV proteins sequence space and drug resistance profiling

The GTMs for PR, RT, and IN inhibitors represent frameworks for visualization of mutant HIV enzymes sequence space and for prediction of their resistance to drugs according to HIVDB thresholds (Figure 2). The maps reflect the density distribution of mutant enzymes that are encoded by different encoding schemes. The color of the maps indicates the mean resistance profile of the sequences residing in a given node of the map. The level of transparency reflects the number of the sequences located in the map. The maps with the highest prediction performance were constructed based on three types of descriptors: 4-mers for PR, one-hot encoded vectors for IN, and PCA-transformed 1-3-mers for RT.

**Figure 2.**
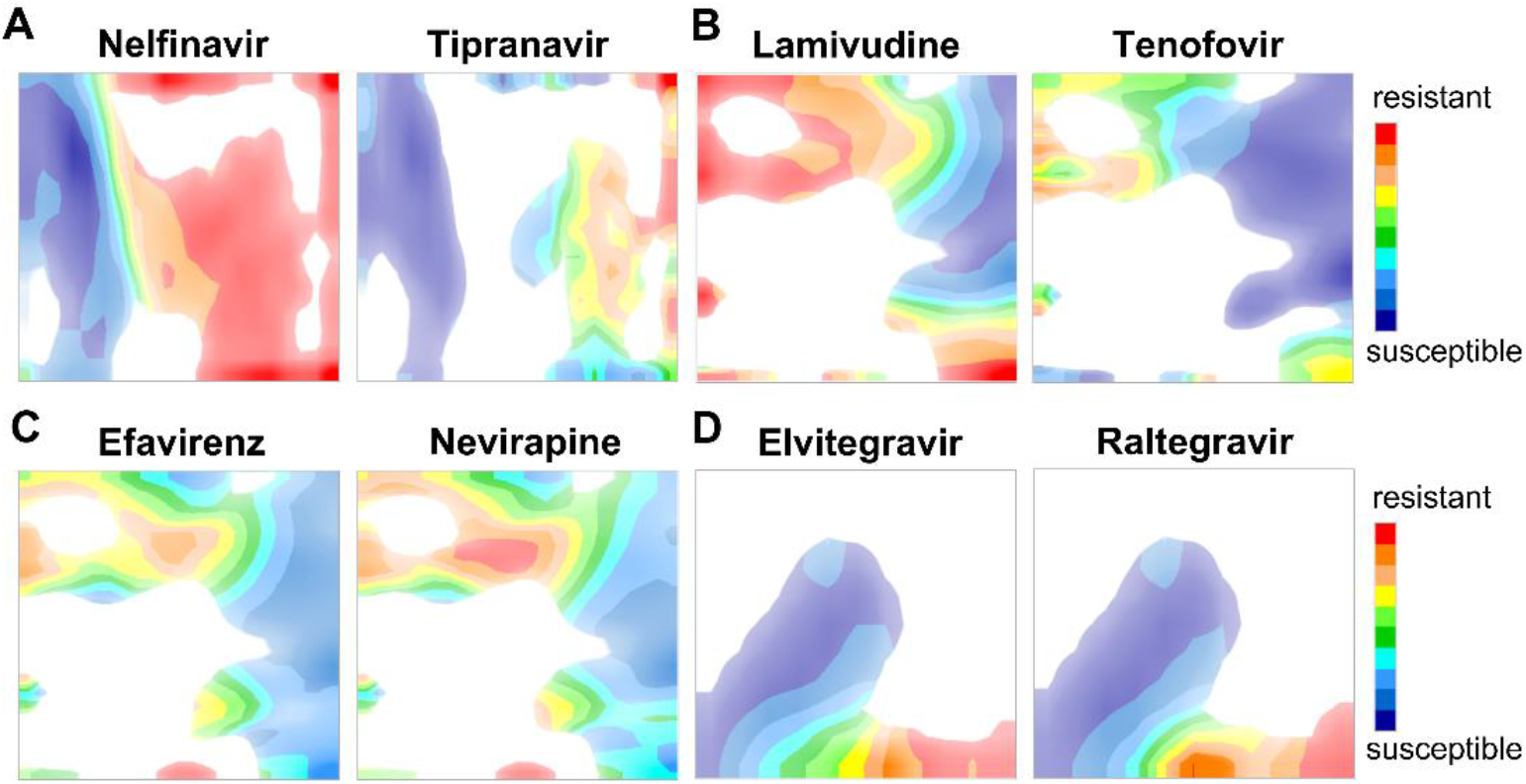
Resistance landscapes for two protease inhibitors (A), two nucleoside reverse transcriptase inhibitors (B), two non-nucleoside reverse transcriptase inhibitors (C), and two integrase inhibitors (D). Each node is coloured by a weighted average of drug resistance profiles of the residing sequences. Red zones are occupied by the resistant sequences, while the blue zones contain susceptible sequences. All colours in between correspond to mixed zones containing both of them. The transparency reflects how many sequences resided in the particular node.

The map for nelfinavir HIV PR inhibitor shows a clear separation between resistant and susceptible PR sequences located on the map (Figure 2A). The comparison of this map to the one for tipranavir reveals common regions that are colored differently, i.e. the same zones on the map are colored in red for nelfinavir, whereas being blue for tipranavir. This highlights the broader spectrum of mutant HIV variants that are susceptible to treatment with tipranavir. Tipranavir is a non-peptidomimetic drug and possesses a different binding profile from other peptidomimetic PR inhibitors. The high potency of tipranavir against even multi-drug resistant HIV PR arises due to a different kind of hydrogen bond patterns that it forms with the residues from the flap region of the PR. Overall, stronger binders such as tipranavir and darunavir are more difficult to be discouraged by mutations.

The resistance landscapes for the nucleoside RT inhibitors were, in general, similar (Figure 2B). In contrast, whereas certain zones on landscapes for nucleoside RT inhibitors are red, the same zones for non-nucleoside RT inhibitors appear in blue. Therefore, non-nucleoside reverse transcriptase inhibitors (e.g. efavirenz, nevirapine) can be effective for HIV variants with mutant RT sequences residing in the bottom right corner of the resistance landscape (Figure 2C). Thus, a simultaneous analysis of maps for nucleoside and non-nucleoside inhibitors could provide an insight to optimal drug combinations for ART regimens.

Integrase inhibitors are usually employed in combination with PR and RT inhibitors in treatment-experienced individuals. The integrase landscapes for the inhibitors considered in this work (raltegravir and elvitegravir) are very similar (Figure 2D). Indeed, raltegravir and elvitegravir possess comparable antiviral activity profiles and the presence of the cross-resistance between these drugs is common (Shimura & Kodama, 2009).

### Comparison to other machine learning methods

Various state-of-the-art machine learning (ML) methods were compared with GTM in terms of prediction accuracies. The GTM-based models allowed achieving high prediction performance in cross-validation (see Figure S4 and Table S2 in Supplementary Data) comparable to the one of other machine learning methods. In more detail, in case of multi-task GTM, the balanced accuracy values ranged from 0.82-0.94 for PR inhibitors (the highest average prediction accuracy among all proteins) to 0.65-0.80 for non-nucleoside RT inhibitors (the lowest average prediction accuracy among all proteins). It should be noted that the GTM was run in a multi-task mode (the same model applies to all inhibitors of a particular viral protein), whereas other approaches were used in a single-task mode, i.e. models were trained for each drug separately. The advantage of the multi-task GTM lies in a versatility of the created maps. The GTM model (map) for a given protein is trained to be able to predict simultaneously the resistance against several inhibitors. Therefore, related resistance landscapes built on this map can be used for comparative analysis of resistance profiles of protein mutants to different drugs. Thus, the GTM implicitly enables an intuitive exploration of the resistance profiles and corresponding complex mutation patterns via resistance and mutation landscapes.

### HIV mutation patterns

The maps can be used for the exploration of HIV mutation patterns and their influence on drug resistance. Since the presence of certain types of mutations defines the resistance profile of the HIV variants, the exploration of the relationship between resistance landscapes and the distribution of sequences with specific mutation patterns on the maps (mutation landscapes) can be used to associate mutation patterns with resistance (Figure 2, Figure 3). In this context, the maps can be used either to visualize the distribution of sequences containing the specific mutation pattern (mutation landscape) and compare it with the resistance landscapes or select a specific part of the map and determine which mutations lead to the resistance profile indicated on the map.

**Figure 3.**
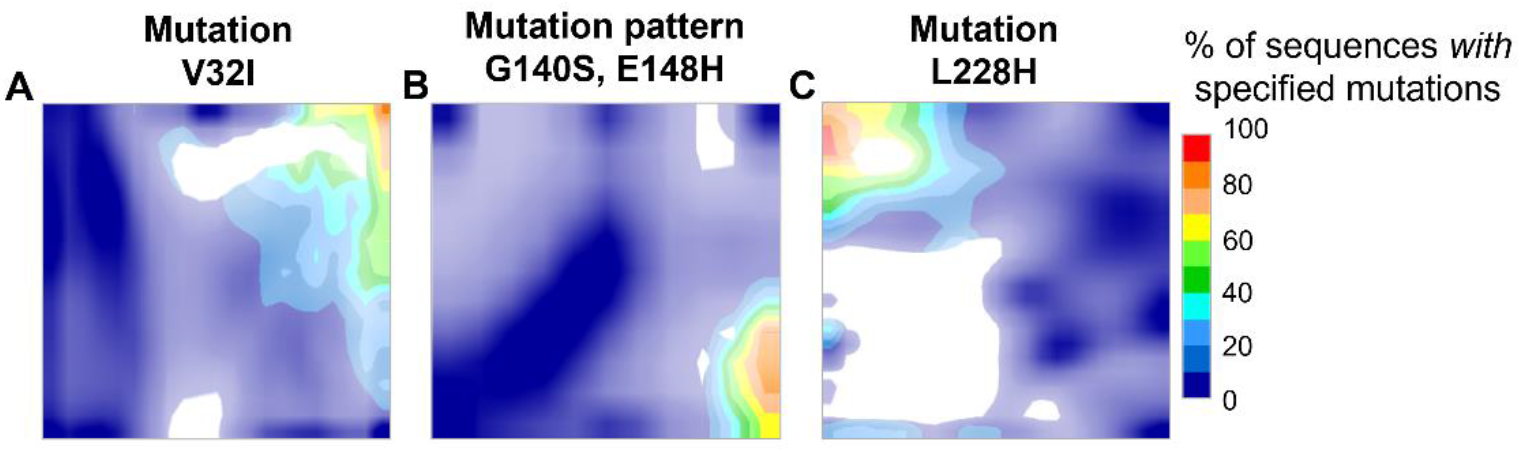
Mutation landscapes for HIV PR (A), IN (B), RT (C) built for 5000 sequences from the frame set. Red zones are predominantly occupied by the sequences with specified mutations, while the blue zones mostly contain sequences without specified mutation pattern. All colors in between correspond to mixed zones containing both sequences with and without specified mutation pattern in different proportions.

At first, the mutation patterns leading to drug resistance for HIV PR, IN, and RT found in the literature (Rhee et al., 2003), (Ceccherini-Silberstein et al., 2007) were analyzed (Figure 2). For example, the key resistance-associated mutation occurring in PR − V32I was investigated (Figure 3). The HIV isolates with this mutation can show a high resistance against all the drugs, including highly potent darunavir and tipranavir (Figure 2 and Figure S5), which can be seen while comparing resistance and mutational landscapes. This is a case of the pan-resistance toward PR inhibitors. All of the PR variants residing in the top right corner of the resistance landscapes are highly mutated. The majority of sequences that contain the V32I mutation also contain other mutations inducing resistance to PR inhibitors. It was experimentally proven that the V32I influences the ability of other mutations to induce resistance against PR inhibitors (Aoki et al., 2018). This effect of the V32I mutation on drug resistance is indeed reflected on the resistance landscapes for all PR inhibitors by the presence of the upper right densely populated red cluster consisting of highly mutated PR sequences containing the V32I mutation among others (Figure 2A, Figure S5). Similarly to the analysis of mutations in protease, both well-known (e.g. G140S, E148H) and emerging resistance-inducing mutation patterns (L228H) in IN and RT, respectively, can also be analyzed (Figure 3). For instance, comparison of the mutation landscapes with the resistance ones (Figures 2–3, Figure S5) shows that these mutation patterns are abundant among mutant proteins conferring resistance.

To identify the residue positions, which are “specific” to the sequences from a particular node or a group of nodes on the map, the SDPred (Kalinina et al., 2009) algorithm was applied (see Methods section). This algorithm, originally developed for prediction of residues responsible for a particular functional specificity of a protein, was repurposed for analysis of predominant mutations that distinguish groups of sequences.

Given the sequences from the selected zone of the GTM, the algorithm was used to predict the set of amino acid alignment positions that are different between the sequences residing in this zone and all other sequences. For example, the sequences located in a zone populated by HIV variants resistant to nelfinavir and susceptible to darunavir were compared to all other sequences (Figure 4). Then, the corresponding specificity determining mutations were established (Table S3). Two mutations with the highest relative frequency of occurrence in the selected zone were established: D30N and N88D (Figure 4A). They are thus the major mutations that differ the sequences from this particular zone and all other sequences residing in the remaining regions of the map. These mutations are indeed associated with strong resistance to nelfinavir according to the literature data (Rhee et al., 2003). Namely, the sequences with the D30N mutation pattern confer strong resistance to nelfinavir, whereas being susceptible to darunavir (Figure 4A). This mutation commonly occurs in combination with 88D inducing cross-resistance to atazanavir and saquinavir. Another mutation L10F is known to induce resistance to both first- and second-generation drugs, namely indinavir, nelfinavir, darunavir, fosamprenavir, and lopinavir, respectively. When the mutations D30N, N88D, L10F occur in a sequence simultaneously (Figure 4B), the resistance profile shifts toward high resistance region for indinavir, lopinavir, saquinavir, fosamprenavir, and atazanavir in comparison to sequences with only D30N, N88D mutation pattern present. Hence, the presence of L10F mutation is important for resistance development. This example illustrates the applicability of the maps for in-depth analysis of mutation pattern-resistance relationships.

**Figure 4.**
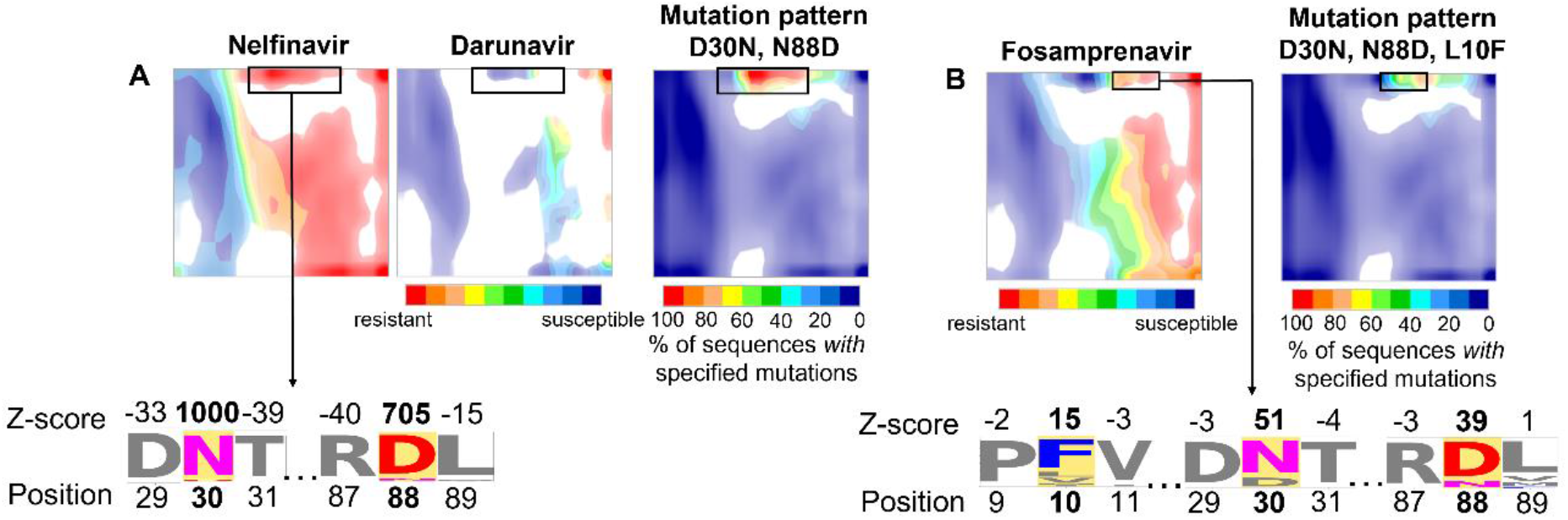
Scheme of the comparative analysis of the resistance and mutational landscapes for HIV PR. (A) The exploration of mutations (D30N, N88D) inducing the resistance to nelfinavir and susceptibility to darunavir from the selected zone of the map (shown as a black rectangle). (B) The exploration of additional mutations (L10F) inducing the resistance to fosamprenavir for the aforementioned selected zone. The coloured stacks of letters highlighted in yellow in the sequence logo at the bottom of the figure represent the amino acids in the specified position in a group of aligned sequences from the selected zone on the map. The coloured letters within the selected stacks represent the resistance determining mutations. The size of the letter reflects the frequency of occurrence of such amino acid in a defined position. The Z-scores (calculated with the SDPred) reflecting which positions within the sequences from the specified zone of the map contain the amino acid mutations that are more prevalent in in sequences from this zone as compared to all others are shown above the corresponding letters.

## Conclusion

To sum up, in this work, a novel cartography-based methodology for protein sequence space exploration and phenotype profiling was suggested. The methodology is based on the non-linear dimensionality reduction method GTM, which allows one to transform complex sequence data and associated phenotypic properties into interpretable two-dimensional maps. While the accuracy of GTM-based drug resistance predictions was comparable with the ones of other state-of-the-art machine learning algorithms, it additionally allows the illustrative analysis of resistance profiles and complex mutation patterns via the resistance and mutation landscapes. The usage of mutation pattern landscapes allows in-depth analysis of complex mutation patterns and leverages the intuitive understanding of their influence on resistance. Hence, this work introduces the first-in-class tool for HIV sequence space exploration and drug resistance analysis combining high predictive accuracy inherent to ML algorithms and interpretability specific to rule-based methods. The introduced methodology is universal and can be applied to other QGPR modelling tasks. Moreover, the GTM can be especially effective for the analysis of big data, since the initial GTM building only requires a limited number of representative samples. Then, the landscapes (QGPR models) can be rapidly built for any amount of available phenotypic data. Since the genomic and proteomic data is rapidly accumulating, the application of the GTM-based methodology for sequences space analysis has the potential to find a wide application as an accurate and interpretable machine-learning framework for personalized medicine.

