## Supplementary Figures and Tables for "HIV-1 drug resistance profiling using amino acid sequence space cartography" (from another version of this preprint: version 1, 2021-08-02)

Table S1. PR, RT, and IN inhibitors with associated fold ratio thresholds from HIVDB (Rhee et al., 2003) used for classification of sequences to drug resistant or susceptible.

| | | Drug | FR(D) |
|---|---|---|---|
| *PR* | *Inhibitors* | Nelfinavir | 3 |
| | | Indinavir | 3 |
| | | Lopinavir | 9 |
| | | Fosamprenavir | 4 |
| | | Saquinavir | 3 |
| | | Atazanavir | 3 |
| | | Darunavir | 10 |
| | | Tipranavir | 2 |
| *Nucleoside RT* | | Abacavir | 3 |
| | | Zidovudine | 3 |
| | | Stavudine | 1.5 |
| | | Didanosine | 1.5 |
| | | Lamivudine | 3 |
| | | Tenofovir | 1.5 |
| *Non-nucleoside RT* | | Efavirenz | 3 |
| | | Etravirine | 3 |
| | | Nevirapine | 3 |
| | | Rilpivirine | 3 |
| *IN* | | Elvitegravir | 4 |
| | | Raltegravir | 4 |

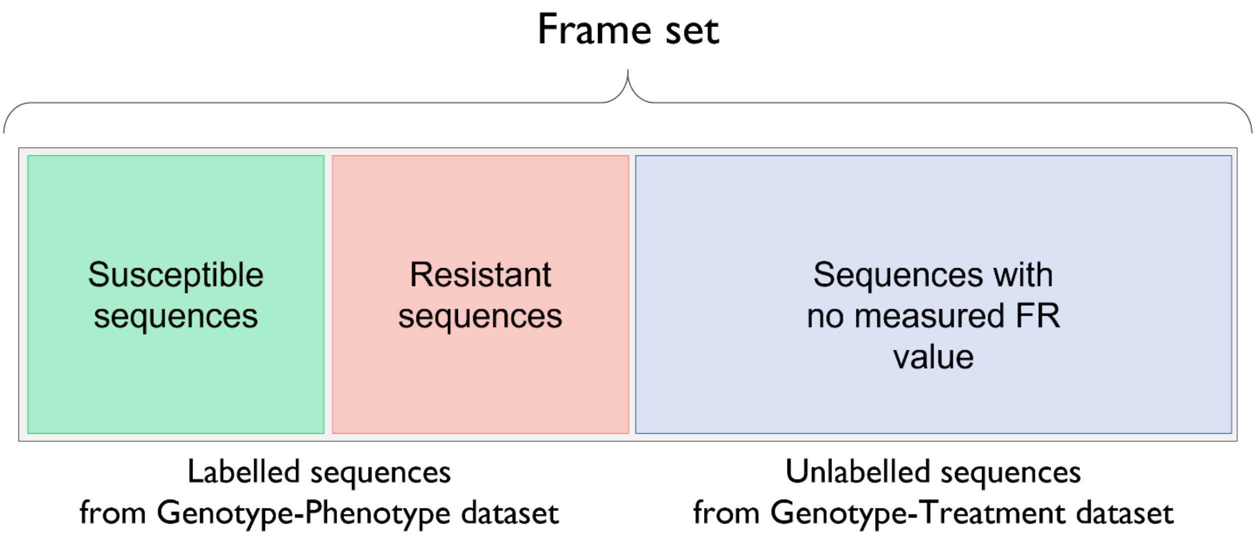


Figure S1. Frame set constitution.

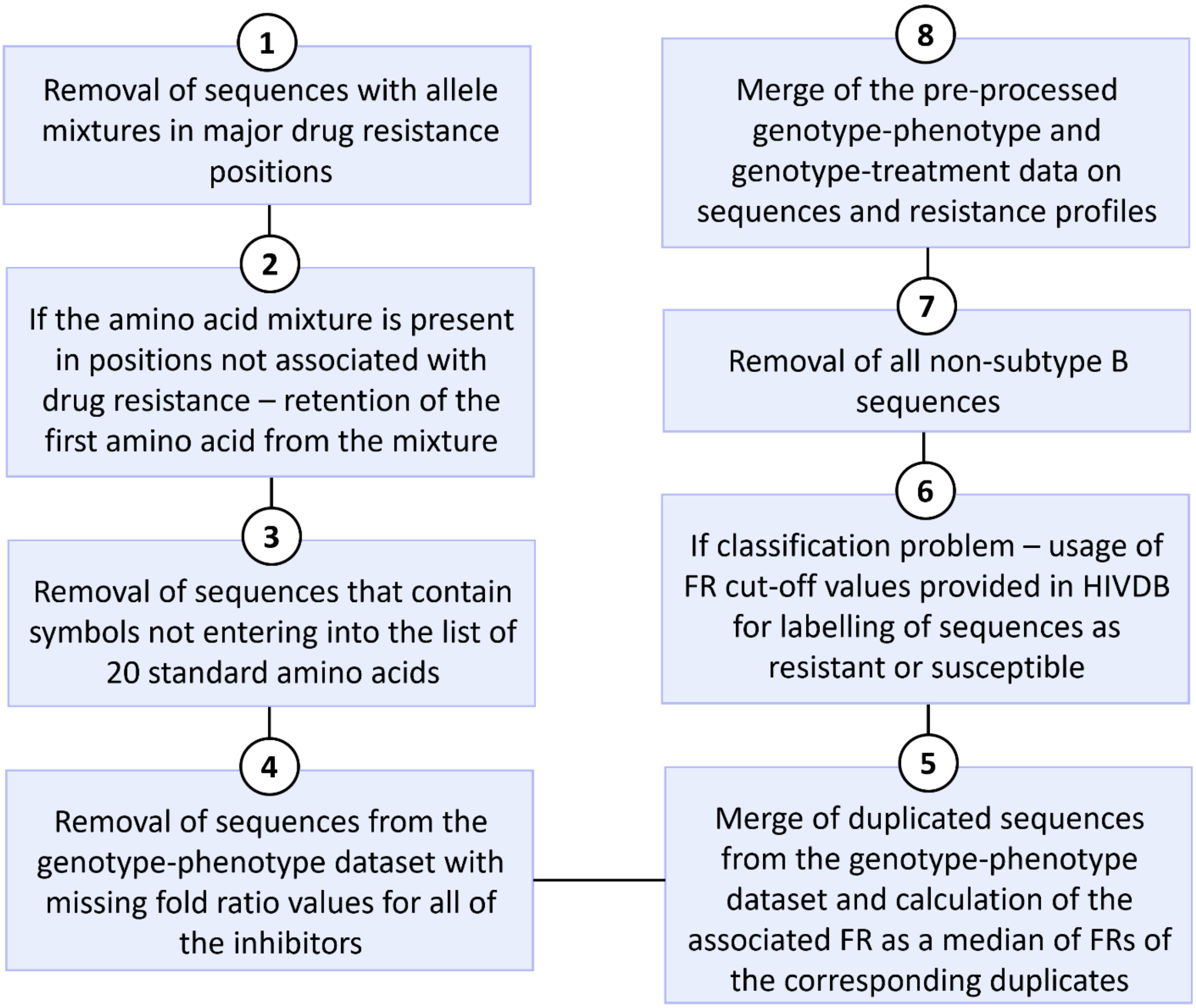


Figure S2. Workflow of data pre-processing that was used in this work.

**Text S3**

The generation of k-mer descriptors ($D_1$, $D_2$, …, $D_N$) consists of counting the number of times a subsequence of a length k (k-mer) occurred in a sequence (Figure S3A). In this context, $D_1$ is the number of occurrence times of a certain k-mer in a sequence, $D_2$ is the number of occurrence times of another k-mer in the same sequence, etc. Hence, a vector of numbers with k-mer counts can be constructed for each amino acid sequence. Descriptors containing k-mers of various lengths taken together as one set of descriptors were also used. Apart from using counts of k-mers, one can employ one-hot encoding scheme consisting of a direct encoding of the whole amino acid sequence that allows conservation of the protein residues' order (Yang et al., 2018) (Figure S3C). In one-hot encoding, each amino acid is represented as a binary vector of length equal to the overall number of considered amino acids. Each amino acid is assigned a number, which corresponds to the index of the bit in the vector that will be assigned a value of '1' with all other bits filled with zeroes. Each of the amino acids from the sequence is encoded with the corresponding binary vector. Subsequently, all of the created binary vectors are concatenated to form a single vector (fingerprint) in accordance with the amino acids' orders in a sequence. In contrast to k-mers, the requirement for the use of one-hot encoding, however is the alignment of all sequences from the dataset, which makes them less adapted for tasks with proteins of various lengths.

While one-hot encoding does not capture any preliminary knowledge on evolutionary relationships among the amino acid sequences (Yang et al., 2018), the BLOck SUbstitution Matrix (BLOSUM) and HIVb matrices allows one to use evolutionary relationships between the sequences (Figure S3D). These matrices contain scores that reflect the frequency of possible exchanges of one amino acid with all others according to aligned sequences from a particular sequence collection (e.g., BLOSUM90 (Henikoff & Henikoff, 1992) – matrix based on aligned sequences of diverse mammalian protein groups that are at least 90% similar, HIVb – matrix based on HIV-1 alignments from different genes encoding viral proteins (Nickle et al., 2007), etc.). In this approach, the vectors derived from the corresponding matrix are used. Once the vectors were assigned to the corresponding amino acids from the sequences, they were concatenated into one single vector similarly to one-hot encoding.

Besides the original descriptors, which can be directly calculated from the sequence, descriptors pre-processed with dimensionality reduction methods, such as PCA and kPCA were used (Figure S3B). The pre-processing can allow one to reduce the time required for manifold building and to obtain smoother landscapes. In this work, both PCA and kPCA methods were applied to all types of descriptors. For kPCA, several pairwise similarity indexes were used as kernels: Hamming distance, Jaccard index, kernels developed by Olier et al. (Olier et al., 2010) based on scores calculated from BLOSUM, and HIVb substitution matrices. In each case, 300 components were preserved and used as the descriptors for the construction of a GTM. In total, 30 descriptors' subsets were generated using an in-house python script ISIDA-Seq.

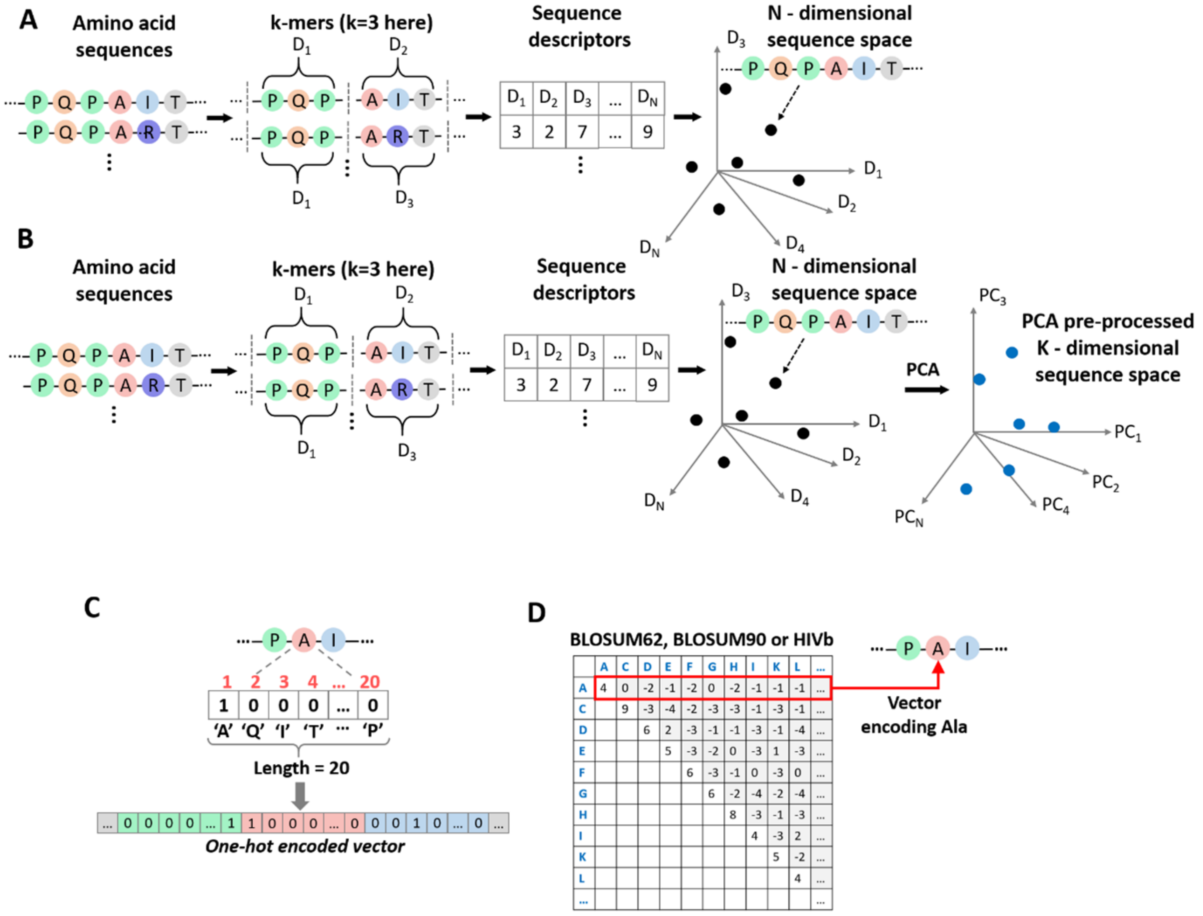


Figure S3. Scheme of sequence descriptors generation: A) k-mers (subsequences of amino acids of length k); B) PCA pre-processed k-mers; C) one-hot encoded vectors; D) Encoding with substitution matrices.

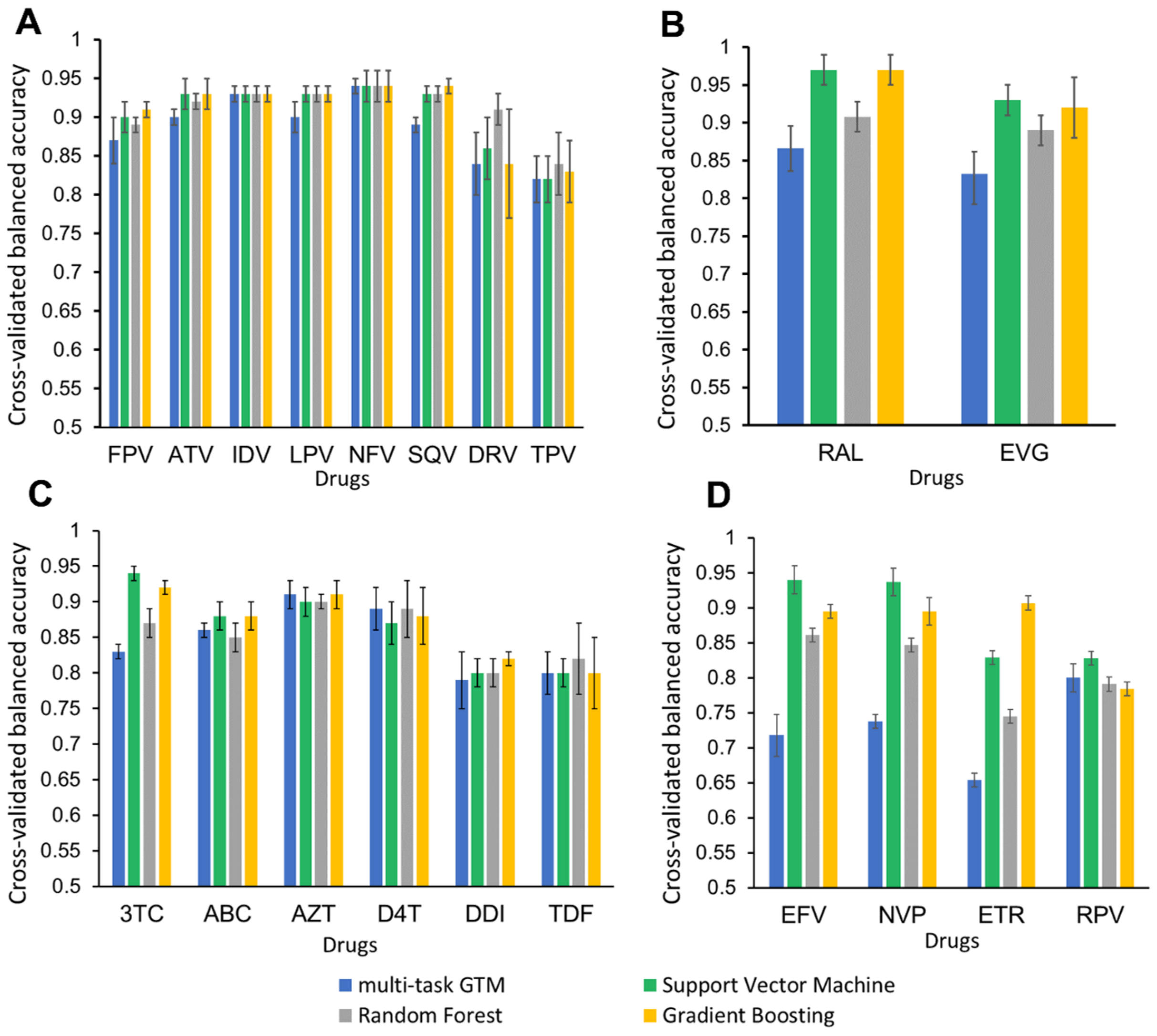


Figure S4. Mean values of BA obtained in 5-fold cross-validation for multi-task GTM (blue), Random Forest (grey), Support Vector Machine (green), Gradient Boosting (yellow) for (A) protease inhibitors: fosamprenavir (FPV), atazanavir (ATV), indinavir (IDV), lopinavir (LPV), nelfinavir (NFV), saquinavir (SQV), darunavir (DRV), and tipranavir (TPV); (B) integrase inhibitors: raltegravir (RAL), elvitegravir (EVG); (C) nucleoside RT inhibitors: lamivudine (3TC), abacavir (ABC), zidovudine (AZT), stavudine (D4T), didanosine (DDI), and tenofovir (TDF); (D) non-nucleoside RT inhibitors: efavirenz (EFV), etravirine (ETR), nevirapine (NVP), rilpivirine (RPV).

Table S2. Mean 5-fold cross validated Balanced Accuracy (BA) values for each of the models build with different methods for each of the anti-HIV drugs.

| | | **Drug** | **5-fold cross-validated Balanced Accuracy (BA)** | | | |
|---|---|---|---|---|---|---|
| | | | Multi-task GTM | RF | SVM | GB |
| *PR* | *Inhibitors* | Nelfinavir | 0.94±0.01 | 0.94±0.02 | 0.94±0.02 | 0.94±0.02 |
| | | Indinavir | 0.93±0.01 | 0.93±0.01 | 0.93±0.01 | 0.93±0.01 |
| | | Lopinavir | 0.90±0.02 | 0.93±0.01 | 0.93±0.01 | 0.93±0.01 |
| | | Fosamprenavir | 0.87±0.03 | 0.89±0.01 | 0.90±0.02 | 0.91±0.01 |
| | | Saquinavir | 0.89±0.01 | 0.93±0.01 | 0.93±0.01 | 0.94±0.01 |
| | | Atazanavir | 0.90±0.01 | 0.92±0.01 | 0.93±0.02 | 0.93±0.02 |
| | | Darunavir | 0.84±0.04 | 0.91±0.02 | 0.86±0.04 | 0.84±0.07 |
| | | Tipranavir | 0.82±0.03 | 0.84±0.04 | 0.82±0.03 | 0.83±0.04 |
| *Nucleoside RT* | | Abacavir | 0.86±0.01 | 0.85±0.02 | 0.88±0.02 | 0.88±0.02 |
| | | Zidovudine | 0.91±0.02 | 0.90±0.01 | 0.90±0.02 | 0.91±0.02 |
| | | Stavudine | 0.89±0.03 | 0.89±0.04 | 0.87±0.03 | 0.88±0.04 |
| | | Didanosine | 0.79±0.04 | 0.80±0.02 | 0.80±0.02 | 0.82±0.01 |
| | | Lamivudine | 0.83±0.01 | 0.87±0.02 | 0.94±0.01 | 0.92±0.01 |
| | | Tenofovir | 0.80±0.03 | 0.82±0.05 | 0.80±0.02 | 0.80±0.05 |
| *Non-nucleoside RT* | | Efavirenz | 0.72±0.02 | 0.86±0.02 | 0.94±0.01 | 0.90±0.01 |
| | | Etravirine | 0.65±0.06 | 0.75±0.03 | 0.83±0.06 | 0.78±0.03 |
| | | Nevirapine | 0.74±0.02 | 0.85±0.03 | 0.94±0.01 | 0.91±0.01 |
| | | Rilpivirine | 0.80±0.09 | 0.79±0.08 | 0.83±0.10 | 0.81±0.06 |
| *IN* | | Elvitegravir | 0.83±0.04 | 0.86±0.02 | 0.93±0.02 | 0.93±0.04 |
| | | Raltegravir | 0.87±0.03 | 0.91±0.02 | 0.97±0.02 | 0.96±0.02 |

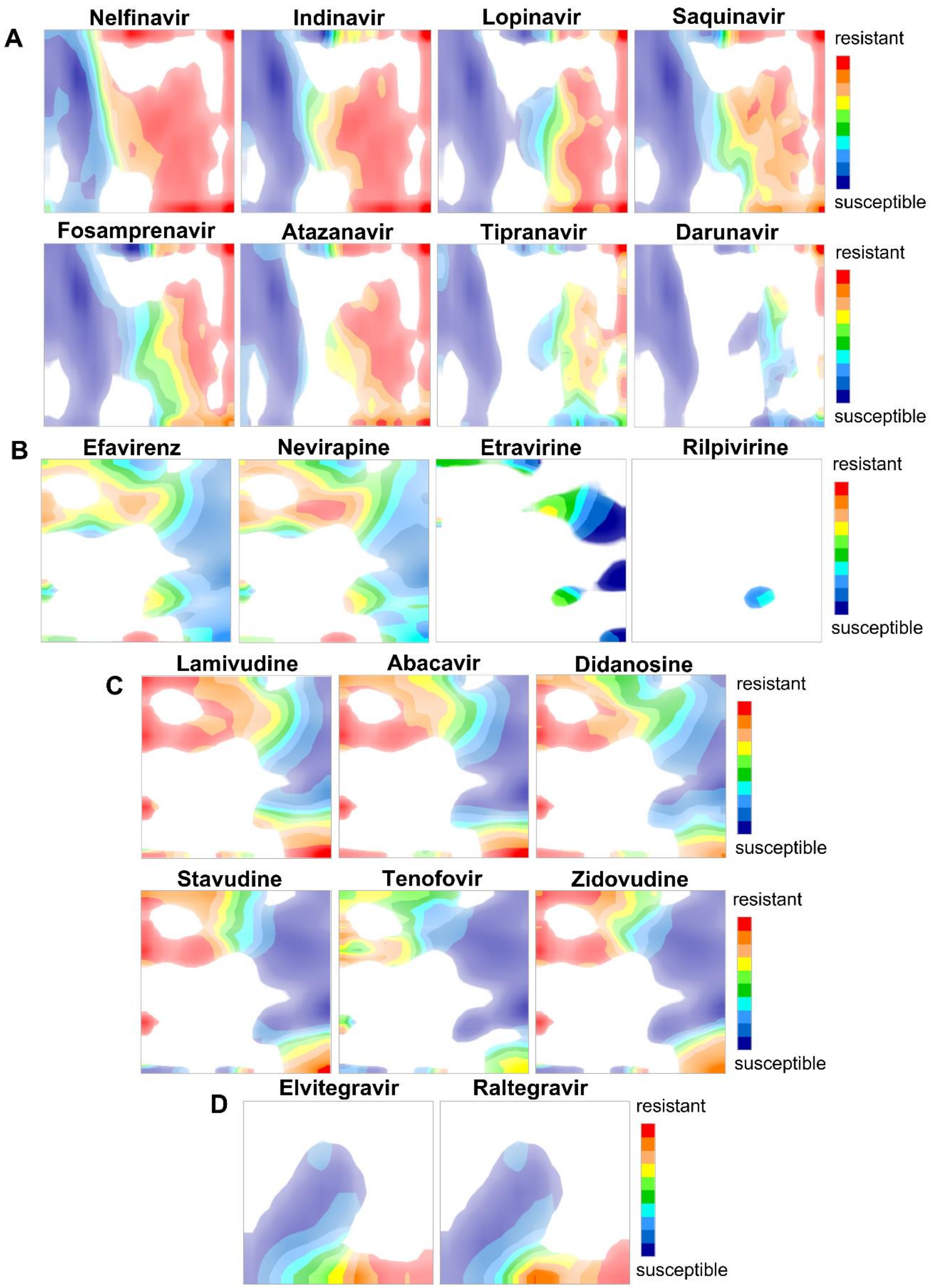


Figure S5. Resistance landscapes for eight protease inhibitors (A), four non-nucleoside reverse transcriptase inhibitors (B), six nucleoside reverse transcriptase inhibitors (C), and integrase inhibitors (D). Each node is colored by a weighted average of drug resistance profiles of the residing sequences. Red zones are occupied by the resistant sequences, while the blue zones contain susceptible sequences. All colors in between correspond to mixed zones containing both of them.

Table S3. The values of Z-score computed with SDPred algorithm for the sequences residing in a zone 1 in comparison to all other sequences; and in a zone 2 in comparison to all other sequences. Zone 1 is populated by HIV variants resistant to nelfinavir and susceptible to darunavir (Figure 4A). Zone 2 is populated by HIV variants resistant to fosamprenavir (Figure 4B).

| Zone | Position | Z-score |
|---|---|---|
| **Zone 1** | 30 | 999.96 |
| | 88 | 704.55 |
| | 10 | 41.85 |
| | 36 | 14.82 |
| | 82 | 14.76 |
| | 63 | 14.47 |
| | 33 | 10.11 |
| | 13 | 8.78 |
| **Zone 2** | 30 | 50.58 |
| | 88 | 38.74 |
| | 33 | 16.39 |
| | 54 | 14.78 |
| | 10 | 14.71 |
| | 36 | 7.11 |
| | 90 | 6.94 |
| | 13 | 6.54 |
| | 20 | 4.34 |
| | 66 | 3.45 |
| | 35 | 2.33 |
